# Syntaxin Habc is required to open Unc18 to template SNARE assembly

**DOI:** 10.1101/2022.05.02.490315

**Authors:** Leonardo A. Parra-Rivas, Mark T. Palfreyman, Thien N. Vu, Erik M. Jorgensen

## Abstract

SNARE and Unc18 proteins form the core of the membrane fusion complex at synapses. The fusion machinery is evolutionarily ancient and mediates constitutive fusion in yeast. We demonstrate that the SNARE and Unc18 machinery in the nematode *C. elegans* can be replaced by yeast proteins and still carry out synaptic transmission. However, substitutions of individual components from yeast disrupts fusion. To understand the functional interactions within the core machinery we adopted an ‘interspecies complementation’ approach using yeast. Synaptic transmission could be restored in chimeras when two key interfaces were present: a novel Habc-Unc18 contact site and an Unc18-SNARE motif contact site. An open form of Unc18 could bypass the requirement for the Habc-Unc18 interface. Together, these data suggest that the Habc domain of syntaxin is required for Unc18 to adopt an open conformation; open Unc18 then templates SNARE complex formation.

## Introduction

In all eukaryotic cells, the fusion of transport vesicles to target membranes requires SNARE and SM (Sec1/Munc18) proteins (Rothman, 2014; Südhof, 2014). For each membrane target, a distinct set of SNARE and SM proteins is used. Fusion at the plasma membrane is mediated by as specific subset of these proteins: In the case of synapses, the SNARE protein on the vesicle is synaptobrevin, the SNAREs on the plasma membrane are syntaxin and SNAP25, and the SM protein is Unc18. The SNARE domains interact at their N-termini and zipper into a four-helix bundle that drives membrane fusion (Gao et al., 2012; Hanson et al., 1997; Min et al., 2013; Pobbati et al., 2006; Sørensen et al., 2006; Sutton et al., 1998; Xu et al., 1999; Zorman et al., 2014).

Syntaxin is composed of 5 domains: an N-terminal peptide, a three-helix bundle called the Habc domain, a linker domain, a SNARE domain, and a transmembrane domain (Fig. S1A). To facilitate trafficking, the synaptic SM protein Unc18 (UNC-18/ Munc18) binds syntaxin in the closed conformation with the Habc domain folded over the SNARE motif (Arunachalam et al., 2008; Han et al., 2009; Hata et al., 1993; Medine et al., 2007; Misura et al., 2000; Rickman et al., 2007; Rowe et al., 2001). At the synapse, SNARE assembly requires syntaxin to be in the open conformation. This transition is thought to be mediated by Unc13 proteins (UNC-13/Munc13) (Gong et al., 2021; Hammarlund et al., 2007; Ma et al., 2011; Magdziarek et al., 2020; Richmond et al., 2001; Yang et al., 2015). Unc18 is required at the final stages to chaperone the SNAREs through their assembly (André et al., 2020; Baker et al., 2015; Gong et al., 2021; Jiao et al., 2018; Lai et al., 2017; Rodkey et al., 2008; Shu et al., 2020; Sitarska et al., 2017). The activation steps between open syntaxin and SNARE complex formation are not fully understood.

To understand the role of the domains of syntaxin in vesicle fusion, we adopted an ‘interspecies complementation’ strategy in the nematode *C. elegans*. Instead of studying mutations generated in syntaxin by random mutagenesis, we substituted entire domains with homologous regions from syntaxins of distant species. In most cases, domain substitutions severely disrupted syntaxin function. The expression of interacting UNC-18 orthologs from the cognate species were then used to restore function. Chimeric versions of UNC-18 further refined interfaces between binding targets.

Notably, replacing the worm Habc domain of syntaxin with the yeast Habc domain severely disrupted neurotransmission. Further replacement of UNC-18 with the yeast homolog Sec1 improved function minimally. Synaptic function was only restored when a chimeric Sec1 could simultaneously interact with both the Habc domain and the SNARE domains. The physiological phenotypes were mirrored by defects in synaptic vesicle docking, demonstrating that morphological docking is a manifestation of SNARE pairing (Imig et al., 2014). Finally, an ‘open’ form of worm Unc18 protein (UNC-18) could partially bypass the requirement for its interaction with its cognate Habc.

Together, the genetic data suggest a model in which UNC-13 signals the presence of a tethered synaptic vesicle and ‘opens’ syntaxin. In the ‘open’ form the Habc domain of syntaxin no longer occludes the SNARE domain. The Habc domain is further required for the transition of UNC-18 from an inactive closed state to an active ‘open’ configuration. UNC-18 in the open conformation binds the SNARE domains of synaptobrevin and syntaxin to template SNARE assembly.

Surprisingly, we also found that substituting the entire yeast SNARE complex along with Sec1 also provided significant rescue – the synapse still functions with yeast proteins, underscoring the conserved functions of these proteins in very different molecular contexts and cellular environments.

## Results

### Syntaxin Habc domain is required for neurotransmission

As a first step toward understanding the function of the domains of syntaxin, we engineered chimeric molecules with the yeast homolog Sso1p. We swapped the N-peptide, the Habc domain, the linker domain, and the SNARE motif (Fig. S1A), and assayed rescue as single-copy transgenes in the null mutant of syntaxin. In *C. elegans,* syntaxin null animals (referred to using the alternative name, *syx-1,* rather than *unc-64,* for clarity) are lethal (Saifee et al., 1998). To study syntaxin null mutants, we rescued animals to adulthood by expressing syntaxin in acetylcholine and glutamate head neurons (Hammarlund et al., 2007). This mosaic approach was used in all instances where the syntaxin transgene did not rescue lethality.

The functions of the chimeric proteins were determined by locomotion assays (Fig. 1A,B) and sensitivity to the acetylcholine–esterase inhibitor aldicarb (Fig. 1C) – resistance to the drug implies a reduced level of acetylcholine release (Mahoney et al., 2006). The N-peptide swap only exhibited subtle changes to locomotion, consistent with some previous experiments (Meijer et al., 2012; Park et al., 2016; Vardar et al., 2021), but not all (Hu et al., 2007; Shen et al., 2007, 2010; Zhou et al., 2013). The replacement of the linker domain resulted in significant defects in locomotion and aldicarb sensitivity. The replacement of the SNARE motif with the yeast Sso1p sequence eliminated syntaxin function in both locomotion and aldicarb sensitivity assays. This is not surprising, given its role in the formation of the SNARE complex. The SNARE motif swap also exhibited reduced syntaxin in axons (Fig. S2A), which is expected since the closed conformation is required for trafficking and the closed conformation requires extensive interactions between the SNARE motif and the Habc domain (Arunachalam et al., 2008; Fan et al., 2007; Han et al., 2009; McEwen and Kaplan, 2008; Medine et al., 2007; Rickman et al., 2007; Rowe et al., 1999, 2001).

**Fig. 1.**
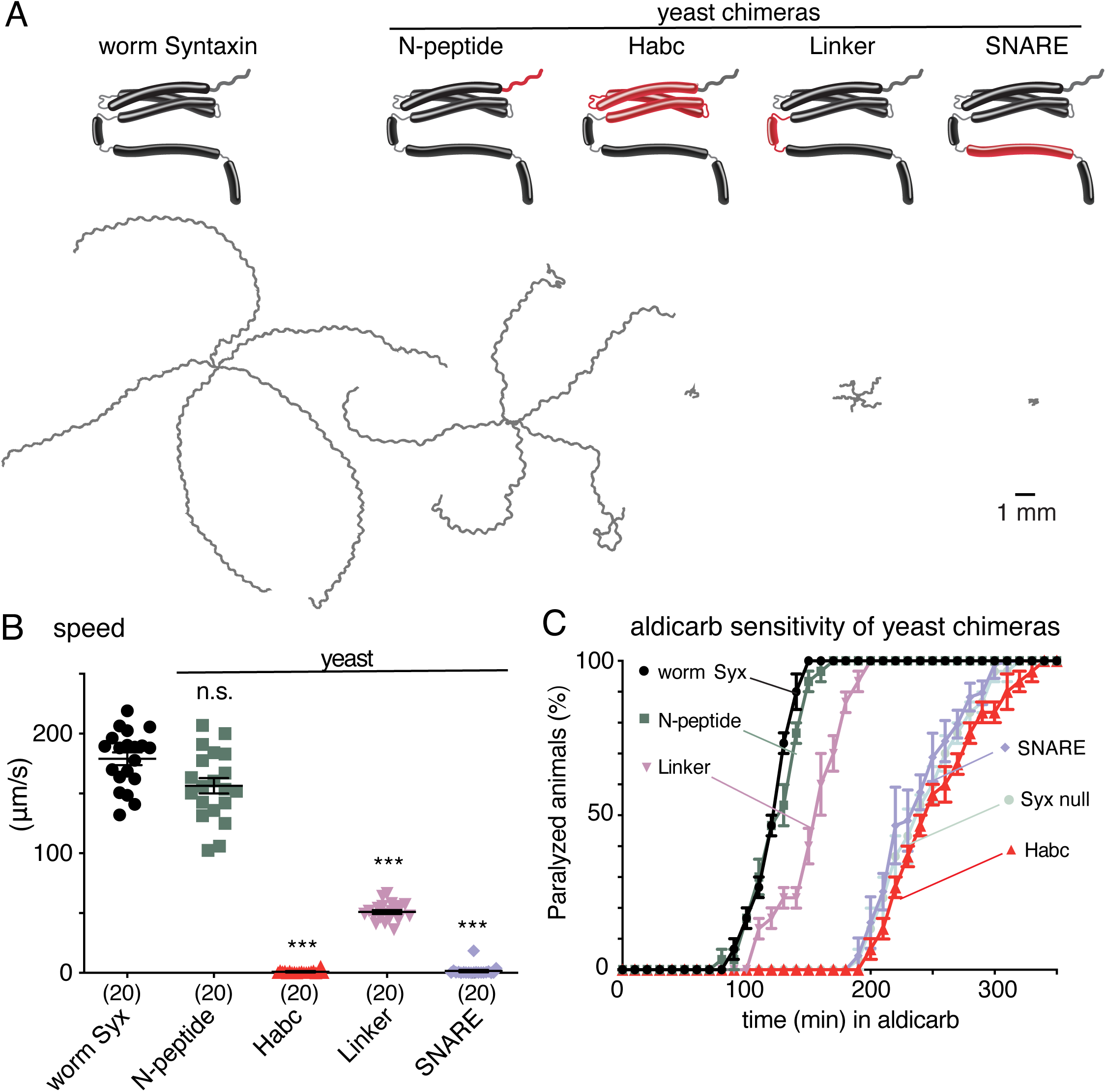
Syntaxin domains differentially contribute to neurotransmission. **A,** (Top) Cartoons depicting syntaxin domains and the chimeras generated by swapping in the corresponding domains from yeast Ssop1. All the chimeras and the rescuing wild-type control (‘worm SYX-1’) are GFP-tagged and integrated into the syntaxin null background, *syx-1.* (Bottom) Representative locomotion trajectories collected for 1 min. **B,** Average locomotion rates (speed) of 20 animals are compared for the same four strains. **C,** Average paralysis time courses after aldicarb exposure (n=3 independent experiments on 20 worms per experiment).

The most intriguing result was that the replacement of the syntaxin Habc domain with the yeast Sso1p Habc domain (yeast-Habc chimera) reduced aldicarb sensitivity and resulted in as severe a defect in locomotion as the SNARE motif swap. The yeast-Habc chimera behaves identically to a full deletion of the Habc domain (Rathore et al., 2010) and to the syntaxin null animals (Fig. 1B-C), demonstrating the importance of this domain. The Habc domain is a conserved, autonomously folding, three-helix bundle (Fernandez et al., 1998), which occludes the SNARE domain, thus preventing the interaction with other SNARE proteins (Dulubova et al., 1999; Misura et al., 2000). This architecture suggests an inhibitory function for the Habc domain. In agreement, the deletion of the Habc domain from Sso1 increases SNARE complex assembly over 2000-fold (Nicholson et al., 1998). However, the deletion of the yeast Vam3 Habc domain (Lürick et al., 2015), the mouse syntaxin Habc domain (Vardar et al., 2021; Zhou et al., 2013), and the worm syntaxin Habc domain (Rathore et al., 2010) all decreased fusion. The decrease in fusion could be attributed to poor trafficking (Fan et al., 2007; Medine et al., 2007; Yang et al., 2006) or poor expression (Vardar et al., 2021; Zhou et al., 2013). We observed some reduced trafficking of syntaxin to axons in our yeast-Habc chimera (Fig. S2A). Again, the Habc-SNARE mismatch should cause syntaxin to adopt the open state. Consistent with this expectation, a constitutively open form of syntaxin (Dulubova et al., 1999) exhibited a similar trafficking defect as the Habc-SNARE mismatch present in the yeast-SNARE chimera and the yeast-Habc chimera (Fig. S2A). However, 70% of syntaxin was properly trafficked and localized to axons in the yeast-Habc chimera.

To determine at what point in evolution the Habc domain acquired characteristics to support synaptic transmission, we swapped in the Habc domains from placozoa, choanoflagellates, and yeast (sequence identities 46%, 38%, and 23% respectively; Fig. S1). The placozoan, *Trichoplax adhaerens*, is a basal multicellular metazoan that possesses the molecular machinery for synapses but lacks neurons and synapses at an anatomical and ultrastructural level (Smith et al., 2014). The choanoflagellate, *Monosiga brevicollis*, is a flagellated eukaryote, which can assume single-celled or colonial forms, and represent a stage prior to the advent of multicellularity (Brunet and King, 2017). By definition, the communication in choanoflagellates takes place between organisms rather than between cells within an organism. The yeast, *Saccharomyces cerevisiae*, is a single-celled organism that communicates to neighbors by the exocytosis of diffusible pheromones (Merlini et al., 2013).

Defects in neurotransmission in nematodes expressing syntaxin Habc chimeras were ascertained by locomotion (Fig. 2A,B), aldicarb sensitivity (Fig. 2C), and electrophysiology (Fig. 2D,E). Trafficking was assayed by fluorescence microscopy (Fig. S2B). All of the syntaxin Habc chimeras were tagged with GFP at the N-terminus. Tagging syntaxin at the N-terminus resulted in a mild reduction in miniature postsynaptic currents (minis/s: wild type: 48.5 ± 7.0; GFP-tagged syntaxin: 33.0 ± 2.7). Surprisingly, both the placozoan and the choanoflagellate chimeras provided a substantial rescue despite a sizable sequence divergence (Fig. S1B). In *Trichoplax*, the rescue was indistinguishable from worm syntaxin (minis/s: worm: 33.0 ± 2.7; *Tricho* Habc chimera: 29.9 ± 3.3). The choanoflagellate chimera provided an intermediate rescue (minis/s: choano Habc chimera: 12.9 ± 2.0). Only the yeast chimera was unable to provide any rescue and was indistinguishable from animals with a full deletion of the Habc domains (Rathore et al., 2010) and from syntaxin null animals (minis/s: *syx-1* null: 0.03 ± 0.02; yeast-Habc chimera: 0.10 ± 0.06). These results argue that the function of the Habc domain must predate synaptic transmission and, to a large extent, even metazoans.

**Fig. 2.**
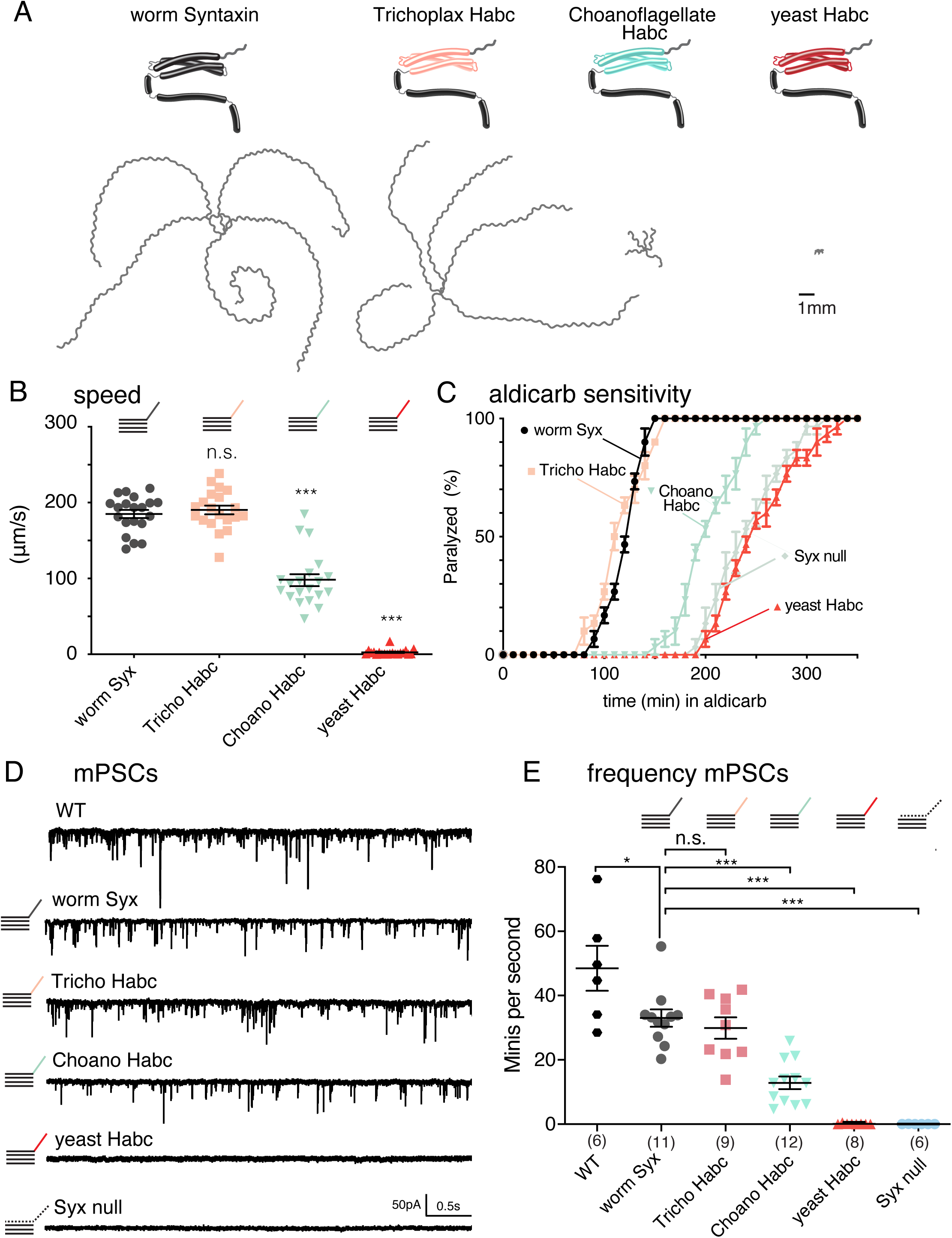
The syntaxin Habc is required for neurotransmission. **A,** (top) Cartoons depicting syntaxin with the Habc domain swapped in from the placozoan *Trichoplax adhaerens*; the choanoflagellate *Monosiga brevicollis*; and yeast *Saccharomyces cerevisiae.* All the chimeras and the wild-type control (worm SYX-1) are GFP-tagged and expressed in the *syx-1* null strain. The GFP tag mildly decreases function compared to the true wild-type (WT) control. (bottom) Representative locomotion trajectories collected for 1 min. **B,** Average locomotion speed of 20 animals compared for the same four strains. **C,** Average paralysis time courses after aldicarb exposure (n=3 independent experiments on 20 worms per experiment). **D,** Representative traces of endogenous miniature postsynaptic currents (minis) recorded from the body muscle of syntaxin chimeras. **E,** Quantification of the mini frequency. Neuronal expression of GFP tagged worm syntaxin rescued mini frequency of syntaxin null animals, but not to wild-type levels (WT, 48.5 ± 7.0 minis/second; n=6 vs. worm SYX-1, 33 ± 2.7 minis/second; n = 11 vs. *syx-1* null, 0.03 ± 0.019 minis/second; n = 6). Mini frequency in Trichoplax Habc chimeras (29.9 ± 3.2 minis/second; n = 9) was not different from rescued worm syntaxin. The average rate of fusion measured from choanoflagellate Habc chimeras (12.9 ± 1.9 minis/second; n = 12) and yeast-Habc chimeras (0.1 ±0.06 minis/second; n = 8) was significantly lower than that measured from the syntaxin rescued strain.

### Habc-Unc18 interactions required for SNARE assembly

The Habc domain in the closed conformation of syntaxin is known to bind Unc18 proteins (Misura et al., 2000). The yeast Unc18 and syntaxin orthologs, Sec1p and Sso1p, bind one another and function in yeast exocytosis (Aalto et al., 1993; Carr et al., 1999; Novick and Schekman, 1979). We reasoned that matching the yeast Habc domain with the cognate Sec1p protein might provide rescue by restoring an interaction interface between these two proteins. In the presence of the yeast-Habc chimera, expression of Sec1p provided a small, but significant, improvement of locomotory function and miniature postsynaptic currents (Fig. 3A,B,C, minis/s: yeast-Habc chimera + Sec1p: 0.13 ± 0.02; yeast-Habc chimera + UNC-18: 0.02 ± 0.00). Similarly, pairing the choanoflagellate Habc chimera with the choanoflagellate SM protein provided significant rescue for locomotion and neurotransmitter release (Fig. S3A,B).

**Fig. 3:**
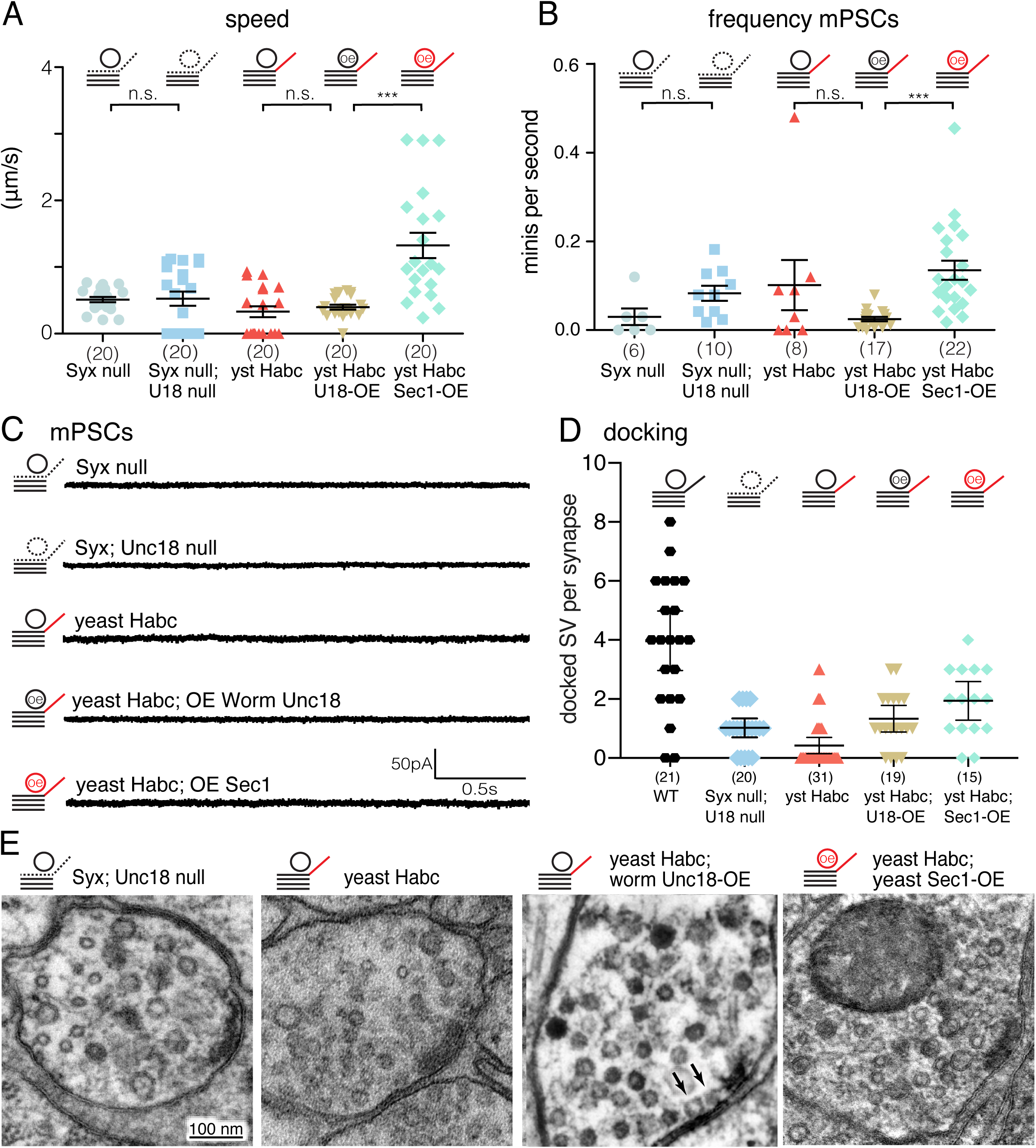
Matching the Habc domain with Unc18 provides only minimal rescue. **A,** Average locomotion rates (speed) in, from left to right: syntaxin null animals; *syx-1 unc-18* double mutants; the yeast-Habc chimera; the yeast-Habc chimera overexpressing worm UNC-18; and the yeast-Habc chimera overexpressing Sec1p (yeast Unc18). **B,** Quantification of the mini frequency in the same five strains, all were similarly defective in synaptic transmission: *syx-1* null: 0.03 ± 0.019 minis/second, n = 6; *syx*-*1 unc-18* null: 0.08 ± 0.017 minis/second, n = 10; yeast-Habc chimera: 0.10 ± 0.057 minis/second, n = 8; yeast-Habc chimera overexpressing worm UNC-18: 0.02 ± 0.005 minis/second, n = 17; yeast-Habc chimera overexpressing Sec1p: 0.13 ± 0.021 minis/second, n = 22. Note, *syx-1* null and yeast-Habc are reproduced from Fig. 2. **C,** Representative traces of endogenous miniature postsynaptic currents (minis) recorded from the body muscle. **D,** Quantification of docked synaptic vesicles at acetylcholine synapses in the *syx-1 unc-18* double mutants, the yeast-Habc chimera, the yeast-Habc chimera overexpressing worm UNC-18, and the yeast-Habc chimera overexpressing Sec1p (yeast UNC-18). All had similarly reduced docking compared to wild-type animals (docked SVs/ per ACh synapse: the wild type, 3.9±0.48, n=21; *syx*-*1 unc-18*, 0.95 ± 0.15 docked SV/synapse, n=20; yeast-Habc chimera, 0.35 ± 0.14, n=31; yeast-Habc chimera overexpressing UNC-18: 1.3 ± 0.21, n=19; yeast-Habc chimera overexpressing Sec1p: 1.9 ± 0.31, n=15). **E,** Representative electron micrographs of the neuromuscular junctions in the ventral nerve cord in the respective strains. For all data, mean and SEM are shown.

Unc18 and syntaxin have both been reported to play a role in synaptic vesicle docking (Hammarlund et al., 2007; Toonen et al., 2006; Voets et al., 2001; Weimer et al., 2003; de Wit et al., 2006). We assayed docked vesicles by reconstructing synaptic regions from serial electron micrographs. The yeast-Habc chimera exhibited 91% and 95% reduction in docking at acetylcholine and GABA synapses, respectively (Fig. 3D,E, docked vesicles: ACh, wild type: 3.9 ± 2.2; yeast-Habc chimera: 0.35 ± 0.14; Fig. S4B,C, docked vesicles: GABA, wild type: 13 ± 3.5; yeast-Habc chimera: 0.7 ± 0.5), similar to *syx*-*1 unc-18* double mutants (Fig. 3D,E) and *syx*-*1* null animals (Hammarlund et al., 2007). Consistent with the electrophysiological recordings, docking was not restored by matching the yeast-Habc chimera with the yeast conspecific Sec1p (Fig. 3D,E, docked vesicles: ACh, yeast-Habc chimera + Sec1p: 1.9 ± 1.2; Fig. S4B,C, docked vesicles: GABA, yeast-Habc chimera + Sec1p: 1.4 ± 0.9). Importantly, the docking defects were not a result of decreased synaptic vesicle numbers (Fig. S4A). We conclude that the binding of Unc18 and the Habc domain is not sufficient to restore synaptic functions: other interactions must be required.

The SM family proteins, which include Unc18, are known to interact with SNARE domains, potentially to template SNARE pairing (Jiao et al., 2018; Lee et al., 2020; Ma et al., 2015; Wang et al., 2019). The crystal structure of the yeast SM protein Vps33 with its cognate SNAREs indicates that the Qa-SNARE and the R-SNARE are bound in what may be a half-zippered SNARE complex (Baker et al., 2015). This structure is likely to have captured an SM protein in the middle of templating the SNAREs. The lack of templating of the synaptic SNAREs by Sec1p could explain the lack of rescue in our Habc-SM match. To test this model, we restored templating to our chimeric proteins. Although yeast Vps33 is only 16% identical to either yeast Sec1p or worm UNC-18, structure predictions and sequence alignments generated by the SWISS-MODEL and Clustal Omega Webservers, respectively (Higgins and Sharp, 1988; Schwede et al., 2003) identified potential residues in worm UNC-18 that would interact with SNARE domains (Fig. S5). We narrowed the list of residues to those that were not conserved between the yeast Sec1p and the worm UNC-18. We replaced these potential SNARE interacting residues on the yeast Sec1p with the corresponding worm residues to generate a ‘Sec1p chimera’ (Fig. S5). Co-expressing the yeast-Habc chimera with the Sec1p chimera yielded a dramatic rescue of synaptic transmission (Fig. 4A,B,C, minis/s yeast-Habc chimera: 0.10 ± 0.06; yeast-Habc chimera + Sec1p chimera: 9.62 ± 1.64). Locomotion and mini rates were improved 60-fold compared to the strain matching only the yeast Habc domain with yeast Sec1p. Thus, when the Unc18 protein is able to interact conspecifically with both the Habc and SNARE motifs, synaptic function is restored (in this case the yeast Habc domain with yeast Sec1p, and the worm UNC-18 SNARE-interacting residues with worm SNARE motifs).

**Fig. 4.**
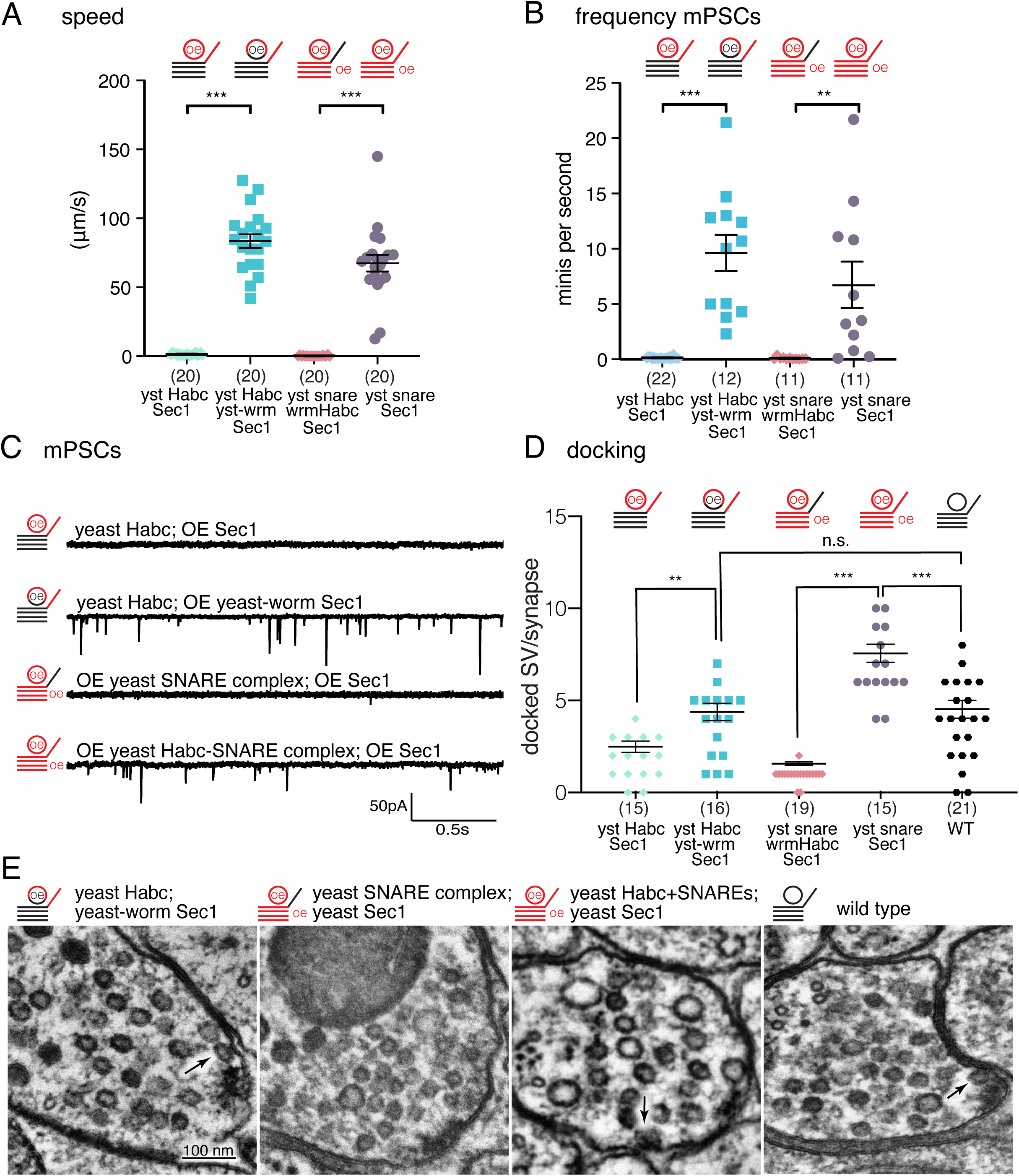
Synaptic transmission is restored with two interaction interfaces: Unc18 – Habc and Unc18 – SNARE domain. **A,** Average locomotion speed in, from left to right: the chimeric yeast-Habc chimera overexpressing Sec1p (yeast Unc18); the yeast-Habc chimera overexpressing the Sec1p chimera (yeast Unc18) with the SNARE interactions restored; syntaxin mutants overexpressing the full yeast SNARE complex and Sec1p without a matching Habc interaction; syntaxin mutants overexpressing the full yeast SNARE complex and Sec1p with a matching Habc interaction. **B,** Quantification of the mini frequency in the same four strains. When the two interaction surfaces are restored, synaptic transmission is rescued: yeast-Habc chimera overexpressing Sec1p: 0.13 ± 0.021 minis/second, n = 22; yeast-Habc chimera overexpressing the Sec1p chimera: 9.62 ± 1.636 minis/second, n = 12; overexpression of yeast SNARE complex with worm Habc + overexpression of Sec1p: 0.09 ± 0.033 minis/second, n = 11; overexpression of yeast Habc-SNARE with yeast Habc + overexpression of Sec1p: 6.70 ± 2.095 minis/second, n = 11). Note, yeast-Habc chimera + overexpression of Sec1p is reproduced from Fig. 3. **C**, Representative traces of endogenous miniature postsynaptic currents (Minis) recorded from the body wall muscle. **D,** Quantification of docked synaptic vesicles in the yeast-Habc chimera overexpressing Sec1p; the yeast-Habc chimera overexpressing Sec1p with the SNARE interactions restored; syntaxin mutants overexpressing the full yeast SNARE complex and yeast Sec1p without a matching Habc interaction; syntaxin mutants overexpressing the full yeast SNARE complex and Sec1p with a matching Habc interaction; and wild-type animals. When the two interaction surfaces are restored synaptic vesicle docking is restored (docked SVs/ per ACh synapse: yeast-Habc chimera + overexpression of Sec1p: 1.9 ± 0.31, n=15; yeast-Habc chimera + overexpression of the Sec1p chimera: 3.8 ± 0.47, n = 16; overexpression of the yeast SNARE complex with worm Habc + overexpression of Sec1p: 0.95 ± 0.093, n = 19; overexpression of the yeast SNARE complex with yeast Habc + overexpression of Sec1p: 6.9 ± 0.49, n = 15; wild type, 3.9±0.48, n=21). Note that the wild type, and the yeast-Habc chimera with overexpression of Sec1p, are the same data as Fig. 3d. **E**, Representative electron micrographs of the neuromuscular junctions in the ventral nerve cord in the respective strains. Arrows indicate docked vesicles.

The dramatic rescue observed with the Sec1p chimera was not due to gain-of-function activity. In the presence of the worm Habc domain, the Sec1p chimera did not confer rescue and was no different than an *unc-18* null mutant (Fig. S6). Note that the *C. elegans* genome encodes a paralog of *unc-18*, T07A9.10, which is expressed ubiquitously, and likely explains the unusually high level of synaptic transmission in *unc-18* null mutants compared to equivalent deletions in other organisms (Cao et al., 2017; Taylor et al., 2021). Thus, the absence of rescue of *unc-18* null mutants, indicates that the Sec1p chimera can only function when it can interact with both the Habc domain and the SNARE motifs.

Docking was also restored when the Sec1p chimera could interact with both the Habc domain and the SNARE motifs (Fig. 4D,E ACh docked vesicles, wild type: 3.9 ± 2.2; yeast-Habc chimera + Sec1p chimera: 3.8 ± 1.9). Interestingly, although the rescue of docking is complete, the rescue of vesicle fusions is only partial (29%) (minis/sec: WT, 33.0 ± 2.7; yeast-Habc chimera + Sec1p chimera, 9.6 ± 1.6) (Fig. 4A-E and Fig. S4B).

In the experiments described so far, the SNARE domains were from the worm. To determine if yeast SNARE motifs can drive fusion at worm synapses, we expressed yeast Sec1p with the entire yeast SNARE complex including the yeast Habc domain (Fig. S7-S8). Remarkably, the yeast Sec1p and the yeast SNARE complex provided substantial rescue – the range was similar to animals co-expressing the yeast-Habc chimera and the Sec1p chimera (Fig. 4A,B,C, minis/s, yeast SNARE complex: 6.70 ± 2.10; yeast-Habc chimera + Sec1p chimera: 9.62 ± 1.64). Importantly, this rescue was completely lost when the Habc domain of yeast syntaxin (Sso1p) was replaced by the Habc domain from worm syntaxin (Fig. 4A,B,C minis/s yeast SNARE complex: 6.70 ± 2.10; yeast SNARE complex with worm Habc: 0.09 ± 0.03). Similarly, docking was rescued by the yeast Sec1p-SNARE complex, but not if the Habc domain was replaced by the worm Habc domain (Fig. 4D,E, ACh wild type: 3.9 ± 2.2; yeast SNARE complex with yeast Habc: 6.9 ± 1.9; yeast SNARE complex with worm Habc: 1.9±1.2). In fact, docking was increased almost 2-fold with yeast machinery compared to the wild type, suggesting that some docked vesicles are fusion-compromised in this strain. Fusion-incompetent vesicles could arise due to inefficient coupling of vesicles to calcium channels via UNC-13, poor pairing with the calcium-sensing machinery, synaptotagmin and complexin, or possibly misassembled SNAREs. It is also possible the yeast machinery is not interacting with the machinery that restricts docking to the active zone of neurons, resulting in ectopically docked vesicles in this strain.

Thus, the Habc domain of syntaxin must interact with UNC-18, and UNC-18 must interact with the SNARE domains to nucleate conspecific SNARE pairing. These experiments further suggest that templating SNARE assembly is a deeply conserved feature in SM proteins: this SNARE-binding interface is functionally conserved from SM proteins used in yeast lysosome fusion to synaptic Unc18 proteins in organisms with nervous systems.

### Syntaxin Habc domain is required to open Unc18

Unc18 proteins can adopt two conformations in crystal structures: a ‘closed’ conformation (Misura et al., 2000) or an ‘open’ conformation (Hu et al., 2011). Specifically, the Unc18 domain 3a transitions from a compact furled loop (closed state) to an extended helical structure (open state) (Hu et al., 2011). A P335A mutation in Unc18 favors the helical extension and increases rates of synaptic vesicle fusion (Han et al., 2014; Munch et al., 2016; Parisotto et al., 2014; Park et al., 2017).

One possible model is that the Habc domain is required to convert UNC-18 into the open state. If true, then the constitutively open form of UNC-18 should bypass the requirement for the Habc interaction to UNC-18. Animals expressing the yeast-Habc chimera are indistinguishable from syntaxin null animals, but expression of the constitutively open form of UNC-18 increased speed 6-fold (Fig. 5A; μm/s yeast-Habc chimera + wild-type UNC-18: 0.43 ± 0.32; yeast-Habc chimera + open-UNC-18: 2.83 ± 1.68). Although the rescue was not complete, open-UNC-18 increased neurotransmitter release as assayed by aldicarb-sensitivity (Fig. 5B) and by electrophysiology (Fig. 5C,D; minis/s yeast-Habc chimera: 0.10 ± 0.06; yeast-Habc chimera + open UNC-18: 1.80 ± 0.70). We also found that open-UNC-18 could bypass the defects seen in the choanoflagellate-Habc chimera (Fig. S3C,D). These data suggest that open state of UNC-18 acts downstream of the Habc interaction with UNC-18.

**Fig. 5.**
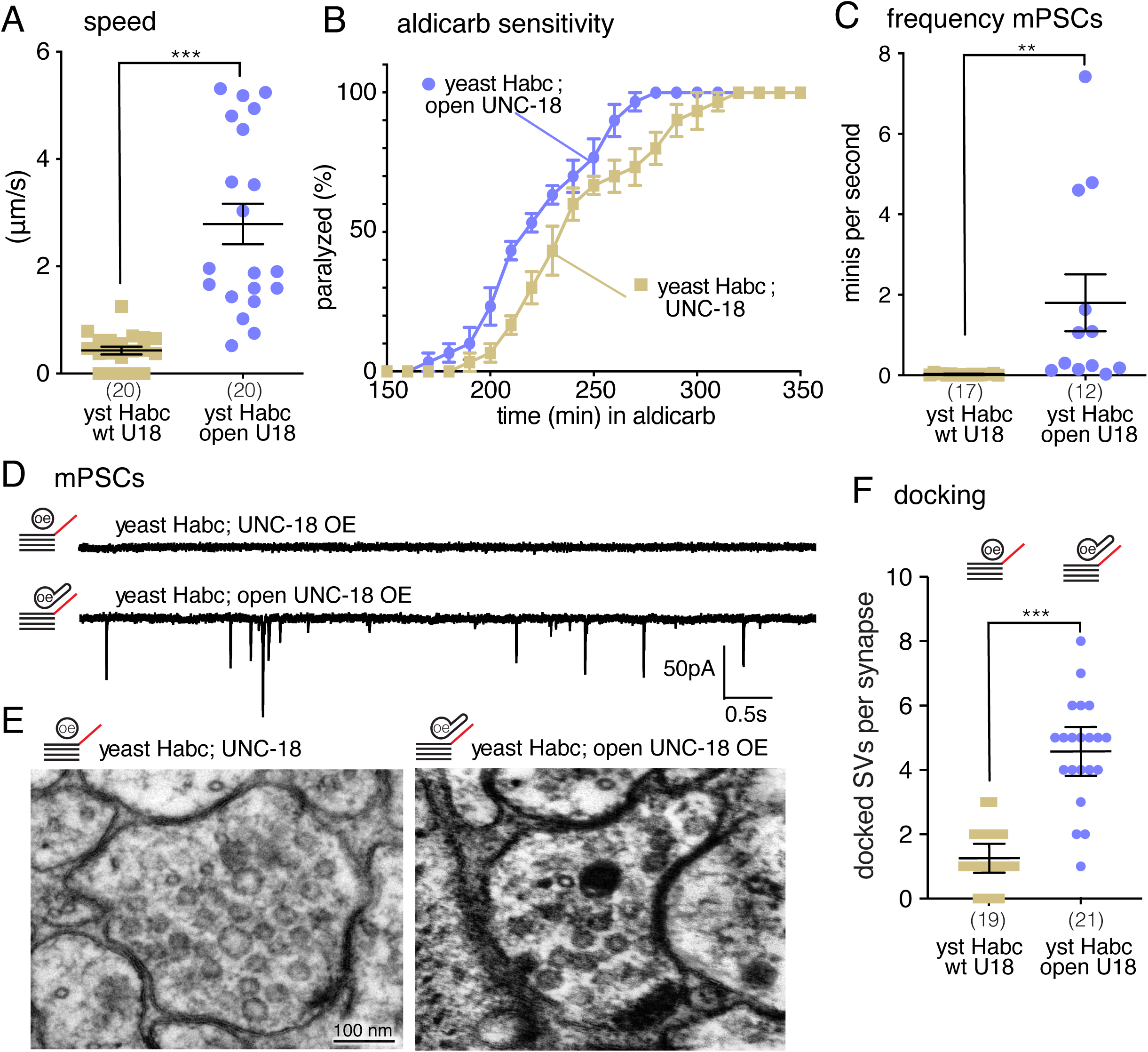
Syntaxin Habc domain opens Unc18. **A,** Expression of P334A UNC-18 mutation ‘Open UNC-18’ in the yeast-Habc chimera background increased the locomotion speed 6-fold (n=20). **B,** Locked open UNC-18 makes yeast-Habc chimeras more sensitive to aldicarb than WT UNC-18 – indicating a restoration of ACh release in the open UNC-18 background. **C,** Open UNC-18 in yeast-Habc chimera increased the frequency of the endogenous miniature postsynaptic currents compare to yeast-Habc chimeras expressing WT UNC-18 (yeast-Habc chimera + overexpression of UNC-18: 0.02 ± 0.005 minis/second, n = 17; yeast-Habc chimera + overexpression of open-UNC-18: 1.80 ± 0.704 minis/second, n = 12). **D,** Representative traces of the endogenous miniature postsynaptic currents from indicated genotypes. **E,** Open UNC-18 restores docking to the yeast-Habc chimeras. Representative electron micrographs of the neuromuscular junctions in the ventral nerve cord in the animals expressing the yeast-Habc chimera with worm UNC-18 (left) or ‘open’ worm UNC-18 (right). **f,** Quantification of docking in the same two strains (yeast-Habc chimera + overexpression of UNC-18: 1.3 ± 0.21 docked SV/synapse, n=19; yeast-Habc chimera + overexpression of open UNC-18: 4.6 ± 0.36 docked SV/synapse, n=21). Note: yeast-Habc chimera + overexpression of UNC-18 data and sample micrograph are the same as Fig. 3d.

To determine if ‘open’ UNC-18 bypassed the requirement for the Habc domain in docking, we performed electron microscopy on the strain expressing open-UNC-18 with the yeast Habc-chimera. The ‘open’ form of UNC-18 bypasses the requirement for the UNC-18-Habc interaction in docking (Fig. 5E,F). Unlike locomotion and physiology, the rescue of docking is complete.

Together, these data suggest a model in which the Habc domain is required to transform UNC-18 from the closed state to the open state, which allows UNC-18 to bind to the SNARE domains and nucleate the SNARE complex formation (Fig. 6).

**Fig. 6.**
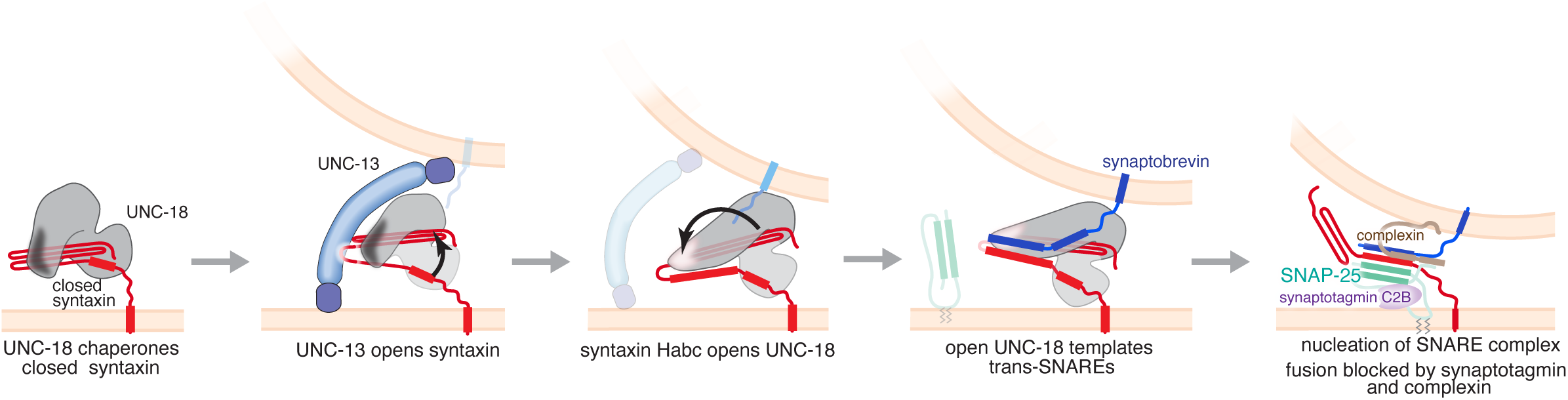
Model of syntaxin Habc domain function. In step 1, Unc18 binds closed syntaxin during trafficking to axons. In step 2, the active zone protein Unc13 converts syntaxin to the open configuration. In step 3, the Habc domain then converts Unc18 to an open conformation. In step 4, open Unc18 binds the SNARE domains of syntaxin and synaptobrevin, to align and nucleate SNARE complex assembly. In step 5, SNAP-25, complexin and synaptotagmin are recruited by unknown mechanisms to form a SNARE complex fully ‘primed’ for fusion. Our results do not explicitly exclude an alternative sequence of steps; for example, the ‘opening’ of UNC-18 could preced the ‘opening’ of syntaxin.

## Discussion

To identify evolutionarily conserved protein interactions in synaptic vesicle fusion, we used an ‘interspecies complementation’ approach to determine the function of the Habc domain in the nematode *C. elegans*. To be most effective, interspecies complementation starts from ‘zero output’, that is, the replacement of a single component from another species resembles a null mutation. We found the worm syntaxin Habc domain could be functionally replaced with those from the Placozoan *Trichoplax* or the Choanoflagellate *Monosiga.* Thus, the conservation of *physical* interactions between the Habc domain and the synaptic machinery predated the evolution of synapses and, indeed, the evolution of metazoans. Eventually, we had to rely on the highly divergent SNARE machinery in yeast to obtain ‘zero output’. The yeast-Habc chimera resembles the syntaxin null mutant and a full deletion of the Habc domain (Rathore et al., 2010).

Fusion was restored by a complex in which two interfaces were species matched: first, the Habc domain and the SM protein, and second the SM protein ‘grooves’ and the SNARE motifs. However, an Habc to UNC-18 mismatch could be bypassed if the UNC-18 protein was locked in the open conformation, thereby suggesting that the Habc domain might function to convert UNC-18 from a closed to open conformation to template SNAREs. Both of these genetic configurations rescued docking, but did not fully rescue fusion rates. One possible reason for incomplete rescue of fusion is that the yeast-Habc syntaxin chimera fails to dock synaptic vesicles adjacent to calcium channels. Unc13 is required for vesicle docking (Hammarlund et al., 2007). In addition, Unc13 couples docked vesicles to calcium channels (Tan et al., 2022) because it binds both Habc domain of syntaxin (Betz et al., 1997) and interacts with calcium channels via RIM (Brockmann et al., 2020; Kaeser et al., 2011). Syntaxin in the open state bypasses the requirement for UNC-13 in vesicle docking, but fusion is only partially restored (Hammarlund et al., 2007). Because of the mismatch between the Habc domain and the SNARE motif, the yeast-Habc syntaxin chimera is predicted to be in the ‘open’ state and dock vesicles independent of Unc13. It is possible that these promiscuously docked vesicles are not coupled to calcium channels, and therefore are not fusion competent. Alternatively, vesicles may be docked near calcium channels but the SNAREs may be misassembled (Lai et al., 2017). Likewise, ‘open’ UNC-18 may bypass crucial steps needed in localizing the fusion machinery near calcium channels or in proofreading SNARE assembly, leading to docked but fusion compromised vesicles.

Two recent structures suggest that the core function of SM proteins is to template SNARE assembly. First, we used the structure of the yeast lysosomal SM protein, Vps33, to map residues in Sec1 that could potentially be modified to interact with worm SNAREs (Baker et al., 2015). Vps33 only has very weak homology with UNC-18, and yet the engineered Sec1 provided significant rescue. The rescue we observed provides the strongest evidence yet for the physiological importance of the templating functions of the SM proteins, which thus far have only been minimally explored in vivo (André et al., 2020). Second, the recent structure of the yeast Golgi SNARE Tlg2 bound to Vps45 indicates that SM proteins can interact with the Habc domain and the SNARE domain in an open conformation (Eisemann et al., 2020). Although similar structural data are not available for Sec1 or Unc18 proteins, binding experiments indicate Unc18 is able to interact with open syntaxin (Christie et al., 2012; Colbert et al., 2013; Rickman et al., 2007; Shen et al., 2007). Finally, our interspecies complementation underscores the universal nature of the interactions between SM proteins and SNAREs that nucleate SNARE assembly.

Taken together our findings suggest a model for the regulatory interactions leading to the SNARE pairing at the synapse (Fig. 6). The active zone protein UNC-13 is thought to convert syntaxin from a closed to an open state (Hammarlund et al., 2007; Ma et al., 2011; Magdziarek et al., 2020; Richmond et al., 2001; Yang et al., 2015), although this role has been disputed (McEwen et al., 2006; Tien et al., 2020). Templating is a late step and requires the open conformation of UNC-18. We therefore speculate that in the open state, the Habc domain of syntaxin converts UNC-18 to an open conformation. Based on the crystal structure of Vps33, the open form of UNC-18 binds the SNARE domains of synaptobrevin and syntaxin to nucleate SNARE assembly (Baker et al., 2015; Sitarska et al., 2017). SNAP-25 is thought to be the last SNARE to enter the complex (Jiao et al., 2018; Kalyana Sundaram et al., 2021; Wang et al., 2019), although the order is still in dispute (Lee et al., 2020).

Perhaps the most remarkable result is that the core fusion machinery from yeast can function in neurotransmission. We do not yet understand how or indeed whether the yeast SNAREs and SM protein are able to couple with the specialized calcium-sensing machinery used in synaptic vesicle fusion. However, the rescue confirms that the functional interactions provided by the SNAREs and their partner SM proteins have remained largely constant from yeast to man, even within very different molecular contexts and cellular environments.

## Acknowledgements

We thank the CGC for maintaining and providing worm strains, and the University of Utah core fluorescence microscopy and electron microscopy facilities for maintenance of equipment used in this study. Britt Graham for suggestions in Methods. Wayne Davis read early versions of the manuscript and provided valuable feedback.

## Contributions

L.A.P. initiated the project with guidance from E.M.J. L.A.P., E.M.J., M.T.P., and T.N.V. designed the experiments. L.A.P. conducted all behavioral, aldicarb, and imaging experiments. M.T.P. conducted electrophysiological experiments. T.N.V. conducted EM experiments. L.A.P., M.T.P., and T.N.V. analyzed the data. L.A.P., E.M.J., M.T.P., and T.N.V. wrote the manuscript.

## Methods

### Strains

All strains were maintained on *E. coli* OP50-seeded NGM plates according to standard methods. Syntaxin null worms are paralyzed and arrest at the first larval stage (L1), which leads to lethality (Saifee et al., 1998). Some of the interspecies chimeras used in our study were unable to rescue *unc-64(js115)* syntaxin null phenotype (referred to using the alternative name *syx-1* in text). To bypass the lethality, we used mosaic animals expressing wild-type syntaxin (Hammarlund et al., 2007) in the acetylcholine neurons of the head; this expression is sufficient to rescue syntaxin null mutants to adulthood. The null allele *unc-18(md299)* is a complete deletion of the locus; the strain is uncoordinated by viable (Weimer et al. 2003). For the *syx-1 unc-18* double mutant we expressed both syntaxin and UNC-18 in the acetylcholine head neurons. These mosaic animals were used in all cases where the syntaxin chimera was unable to rescue lethality (see Table S1 for a complete list of strains used in this work).

### Molecular Biology

All plasmids were made using the Invitrogen multisite Gateway cloning technique. To build the rescuing construct for *unc-64(js115)* (syntaxin null animals), the neuronal UNC-64A cDNA was amplified from a worm cDNA library and cloned into Gateway entry vectors. This parental construct was used to engineer all syntaxin chimeras in this study. Domain replacement was performed by insertion of the corresponding synthetic gene fragments (Integrated DNA Technologies (IDT)) and assembled by Gibson cloning. The resulting [1-2] entry clones were recombined with a [4–1] entry vector containing the synaptotagmin promoter and a GFP tag (pEGB348); a [2–3] entry vector with the let-858 3ʹUTR; and the [4– 3] destination vector (pCFJ201) using LR clonase (Invitrogen). Plasmids were inserted into the worm genome as a single copy using the MosSCI technique (Frøkjaer-Jensen et al., 2008) or expressed as extrachromosomal arrays for overexpression experiments. Similarly, UNC-18 rescuing constructs were obtained by cloning the UNC-18 cDNA or the synthetic Sec1p cDNA (from Biomatik) into the Gateway entry vectors. These plasmids were also used to generate the open UNC-18 mutant and the Sec1p chimera. In both cases, synthetic gene fragments with the desired nucleotide modification were cloned using Gibson assembly. The open UNC-18 mutant was obtained by mutation of the “hinge” proline (P334A) in UNC-18 domain 3a. To build the Sec1p chimera we introduced worm residues in the groove and cleft domains of Sec1p. To identify these substitutions we used the crystal structure of the yeast SM protein Vps33 with its cognate SNAREs (Baker et al., 2015). Residues from the Vps33 predicted to binds Nyv1 and Vam3 (PDB code 5BV0 and PDB code 5BUZ, respectively) were mapped into Sec1p based on secondary structure predictions generated by the SWISS-MODEL Webserver (Schwede et al., 2003) and sequence alignments obtained with Clustal Omega (Higgins and Sharp, 1988) (S5). Using a pymol script we identified potential contact residues in Sec1p within 4 Angstrom that are not conserved between yeast and worm.

### Imaging

Nematodes were immobilized using 25 mM sodium azide (NaN_3_) and mounted on 3% agarose pads on glass slides. All images were acquired as Z-stacks using a Pascal LSM5 confocal microscope (Carl Zeiss) with a 63x 1.4NA oil objective. Ventral cord images were taken with the cord facing toward the objective. Fluorescent intensity was quantified using Image J software. Axon intensity was obtained by drawing a region of interest around the ventral nerve cord including the soma (total intensity) and subtracting the soma intensity. Data was analyzed using one-way ANOVA followed by Bonferroni’s post-test and reported as mean ± SEM.

### Worm tracking and speed analysis

To compare worm tracks, a single young adult worm was placed on a NGM plate seeded with OP50. After 1 minute, the animal was removed and a track picture was taken with a Stingray camera (Allied Vision Technologies model). Worm tracks were then drawn on a WACOM touchscreen monitor, x,y coordinates, and length measurements were determined using an ImageJ macro. ImageJ x,y coordinates were transformed into a scalable vector graphics file (svg) using a Matlab script developed in the Jorgensen lab. In Fig. S8, we allowed the animals to move for 5 minutes, instead of the customary 1 minute, to better distinguish between animals with severe locomotion defects.

To measure the speed, 20 animals for each strain were filmed for 2 minutes. Animals with severe locomotion defects were filmed for 30 minutes. Videos were generated using Wormtracker system (MBF Bioscience). Videos were then analyzed and average speed computed using WormLab software built in with the tracker system.

### Aldicarb assays

Aldicarb sensitivity was assessed using 20 young adult worms on NMG plates containing 2 mM aldicarb. Worms were scored for paralysis at 10 minutes intervals for 6 hours. Worms were considered paralyzed when there was no movement in response to three taps to the head and tail with a platinum wire. Once paralyzed, worms were removed from the plate. Each genotype was tested blind three times and paralysis curves were generated by averaging paralysis time courses for each plate.

### Electrophysiology

Electrophysiological recordings were performed as previously described (Richmond and Jorgensen, 1999; Richmond et al., 1999). Worms were immobilized with cyanoacrylate glue (Aesculap Histoacryl; BBraun Inc.) and a lateral incision was made to expose the ventral medial body muscles. The preparation was treated with collagenase (type IV; Sigma-Aldrich) for 15 seconds at a concentration of 0.5 mg/mL. The muscle was voltage-clamped using the whole-cell configuration at a holding potential of −60 mV. All recordings were performed at 21°C using an EPC-9 patch clamp amplifier (HEKA), which runs on an ITC-16 interface (HEKA). Data were acquired using Pulse software (HEKA). Data analysis and graph preparation were performed using Pulsefit (HEKA), Mini Analysis (Synaptosoft), and Igor Pro (Wavemetrics). Bar graph data are presented as the mean ± standard error of the mean.

### Electron microscopy

Electron microscopy were performed as previously described (Watanabe et al., 2013). Ten young adult worms were placed into a 100-um deep specimen carrier (type A and B) along with space-filling 5% bovine serum albumin in M9 buffer. Samples were frozen with a Leica EM-ICE high-pressure freezer. Freeze substitution was performed in a Leica EM AFS2 system. Frozen samples were fixed with 1% glutaraldehyde, 1% osmium tetroxide, and 1% water in anhydrous acetone for 24 hours at -90°C. The samples were warmed to -20°C at 5°C/hour, held at -20°C for 16 hours, and subsequently brought to room temperature (20°C, at 10°C/hour). Fixed animals were isolated from the specimen carrier and embedded in Epon-Araldite resin. Two random animals from each genotype were sectioned. ∼250 ultrathin (33-40 nm) serial sections for each animal were collected on a Leica microtome (Ultracut UC7). The sections were post-stained with 2.5% uranyl acetate in 70% methanol for five minutes before imaging. Serial micrographs of the ventral nerve cords were collected in a transmission-mode scanning electron microscope (Zeiss GeminiSEM 300). ATLAS 5 was used to acquire images in a semi-automated fashion.

Synaptic morphometry was performed blind to genotype. Micrographs from serial sections encompassing a single synapse were collected as an image stack. A synapse was defined as all profiles containing the presynaptic dense projection plus a flanking profile from each side of the dense projection (Watanabe et al., 2013). If more than two synaptic profiles were missing (section loss, occluding schmutz, etc.), the synaptic series was excluded from the analysis. The stacks of serial synaptic profiles were then randomized. Stacks were then segmented for features, such as total synaptic vesicle numbers, and docked vesicles. A synaptic vesicle was considered docked if the vesicle membrane touches the plasma membrane (0 nm) without lighter pixels between the vesicle and plasma membranes (Hammarlund et al., 2007). Acetylcholine and GABA synapses were segmented based on the *C. elegans* the reconstruction of the connectome from serial electron micrographs (White et al., 1986).

### Multiple Sequence Alignment Analysis and Model Interpretation

Protein sequences were retrieved from Uniprot via Jalview’s sequence fetcher (Waterhouse et al., 2009) and aligned with the Clustal Omega Webserver with default parameters (Higgins and Sharp, 1988). For the syntaxin alignment shown in S1, mouse (uniprot O35526), worm (uniprot O16000-2), *Trichoplax* (uniprot B3S4L5), *Monosiga* (uniprot A9UTG5), and yeast (uniprot P32867) sequences were used. Alignment in S5 used the Vps33 sequence from *C. thermophilum* (uniprot G0SCM5) and UNC-18 sequences from mouse (uniprot A2ARS2), worm (wormbase UNC-18a 591aa, Note: the uniprot sequence P34815 673aa, is an anomalous sequence that appends 91 N-terminal amino acids from a region 5’ of the *unc-18* gene), *Trichoplax* (uniprot B3RPC7), *Monosiga* (uniprot A9V0L3), and yeast (uniprot P30619). Alignment in S7 for synaptobrevin used sequences from worm (uniprot O02495) and yeast (uniprot P1109). Similarly, SNAP-25 alignment used sequences from worm (uniprot A5PEW5) and yeast (uniprot P40357).

All figures containing models (S1B and S5) were prepared with Chimera and ChimeraX (Pettersen et al., 2004, 2021).

### Statistical analysis

Data in scatter plot graphs present single observations (points) and are shown as mean ± standard error of the mean (SEM). Data from two groups with normal distribution or non-parametric distribution were subjected to Student’s two-tailed t-test or Mann-Whitney non-parametric test, respectively. All the tests were performed using GraphPad Prism 8.3. A level of *P* < 0.05 was considered significant.

---

**Fig. S1.**
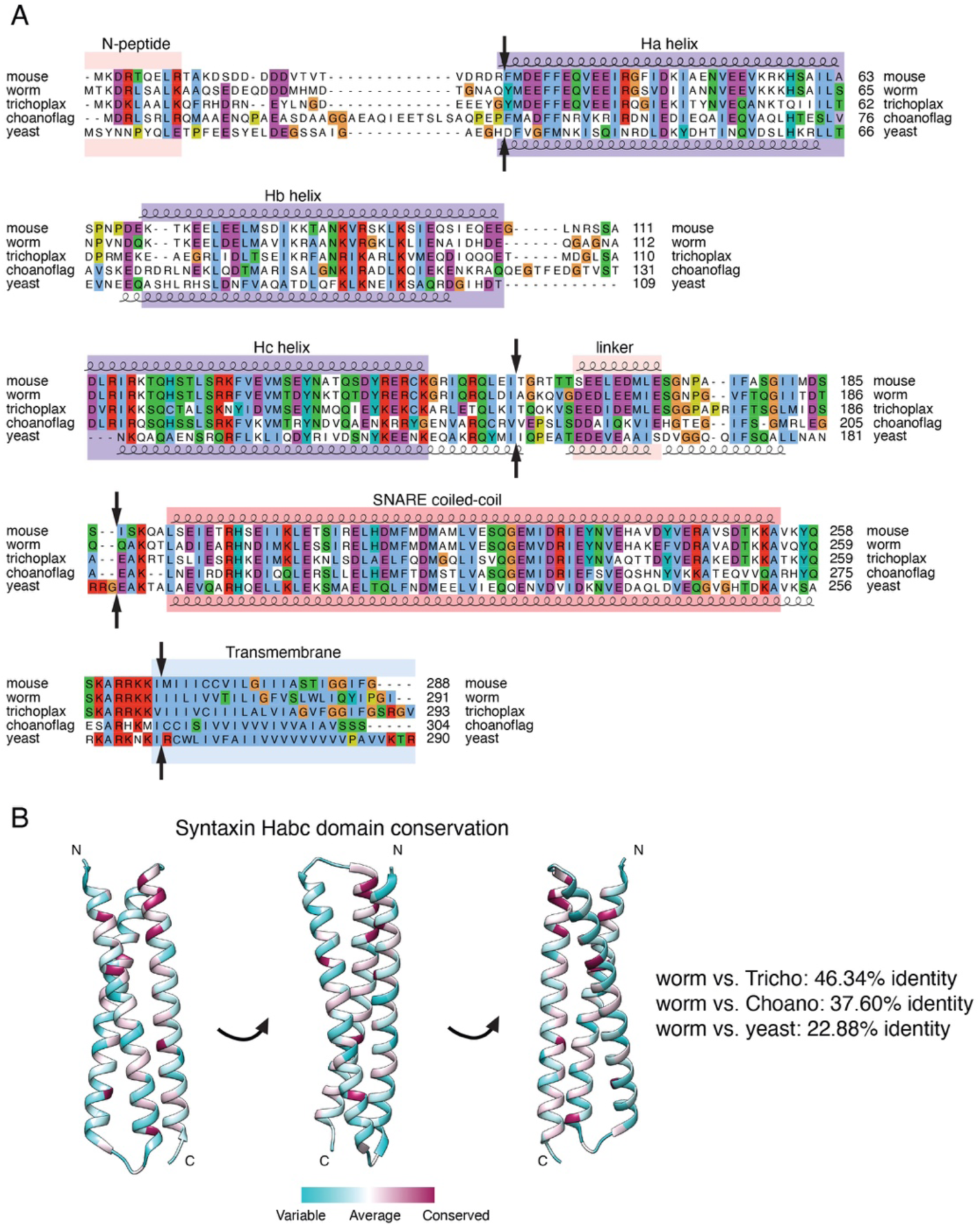
Sequence alignment of syntaxin proteins. **A,** Domain architecture and homology of the syntaxin proteins. Domains are illustrated in blocks above the protein alignment. The black arrows indicate the breakpoints in the chimeric proteins. The alignment uses the default Clustal X color scheme (Larkin et al., 2007): hydrophobic blue, positive charge red, negative charge magenta, polar green, cysteines pink, glycines orange, proline yellow, aromatic cyan, unconserved white. **B,** Relative conservation of the syntaxin Habc domain is based on the species shown in S1A. Habc domain sequences from placozoa (*Trichoplax adhaerens*; Syntaxin 1.2; uniport B3S4L5, choanoflagellates (*Monosiga brevicollis*; predicted protein; uniprot A9UTG5), and yeast (*Saccharomycies cerevisiae*; Sso1p; uniprot P32867) have distinctive amino acid identities compared to the worm Habc domain (*Caenorhabditis elegans*; *unc-64*; uniprot O16000-2).

**Fig. S2.**
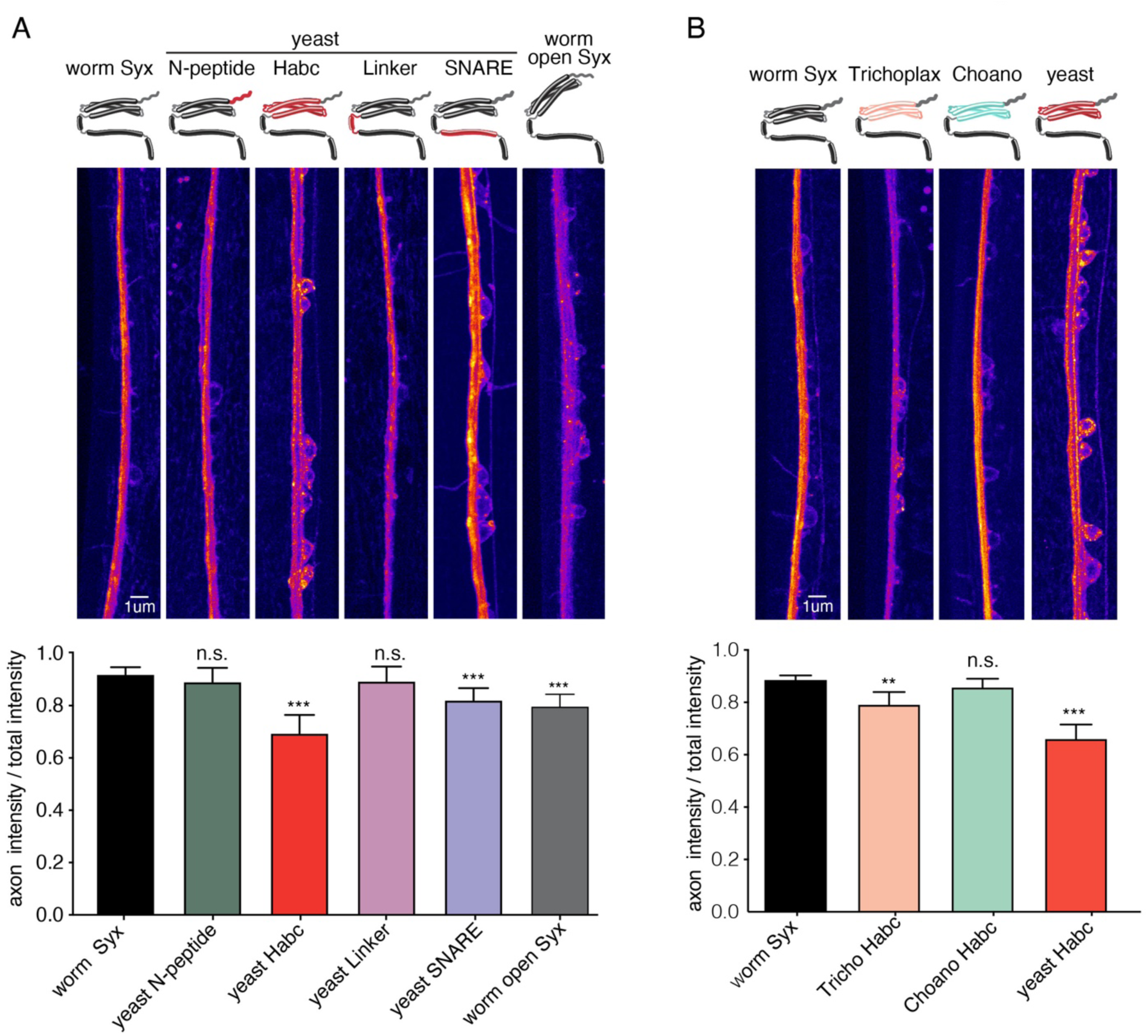
The Habc domain and SNARE domain are important for syntaxin localization. **A,** Substitution of the Habc or SNARE motif of syntaxin disrupts syntaxin localization. **A** (top) Representative images showing the localization of syntaxin chimeras and open syntaxin. The yeast-Habc chimera, the yeast SNARE chimera and syntaxin in the open configuration showed a reduced axonal localization, indicating defects in transport. However, the majority of syntaxin was properly trafficked and localized to axons. Chimeras with the yeast N-peptide and the yeast syntaxin linker had normal syntaxin localization. **A,** (bottom) Quantification of the same five strains used in top panel. **B,** Trichoplax and yeast-Habc chimeras are mislocalized. Substitution of the worm Habc domain with either Trichoplax or yeast Habc impaired syntaxin localization. **B,** (top) Representative images showing the localization of the syntaxin Habc chimeras. Worm syntaxin and the choanoflagellate Habc chimeras were correctly localized to axons with approximately 10% accumulation in neuronal cell bodies. By contrast, Trichoplax and yeast-Habc chimeras exhibited increased localization in cell bodies (20% and 30% respectively). **B,** (bottom) Quantification of the same four strains shown in top panel.

**Fig. S3:**
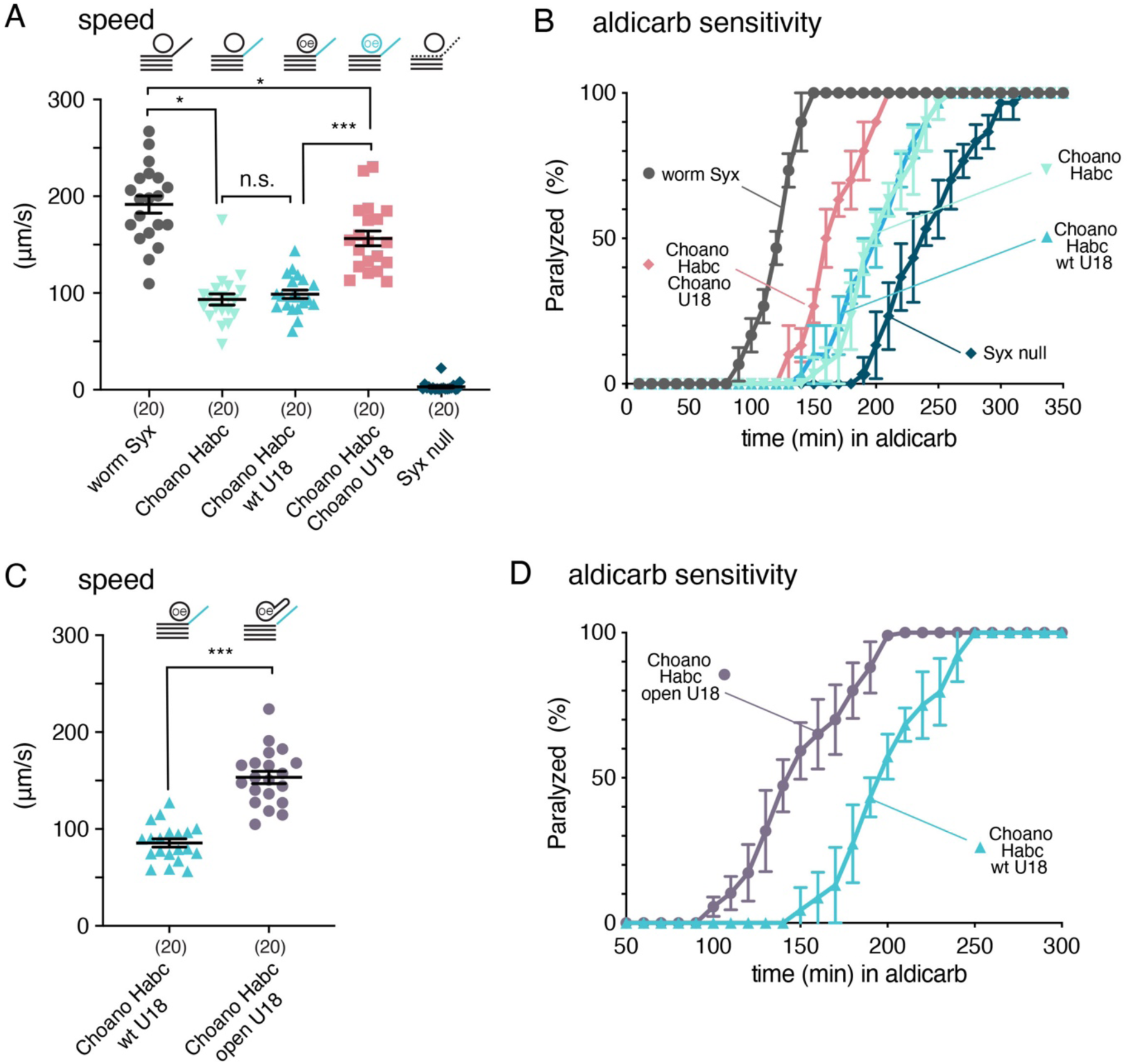
Matching the choano Habc chimera with either its cognate SM protein or an open form of UNC-18 restores neurotransmission. **A,** Average locomotion rates in, from left to right: rescued worm syntaxin; the Choano Habc chimera; the Choano Habc chimera overexpressing worm UNC-18; the Choano Habc chimera overexpressing Choano Unc18; and the syntaxin null animals. **B,** Matching the Habc domain with Unc18 from choanoflagellates provides partial rescue in the aldicarb-sensitivity assay (n=3 independent experiments on 20 worms per experiment). **C,** Expression of P334A UNC-18 mutation ‘Open UNC-18’ in the Choano-Habc chimera background increased the locomotion speed 1.8-fold (n=20). **D,** Locked open UNC-18 makes Choano-Habc chimeras more sensitive to aldicarb than WT UNC-18.

**Fig. S4.**
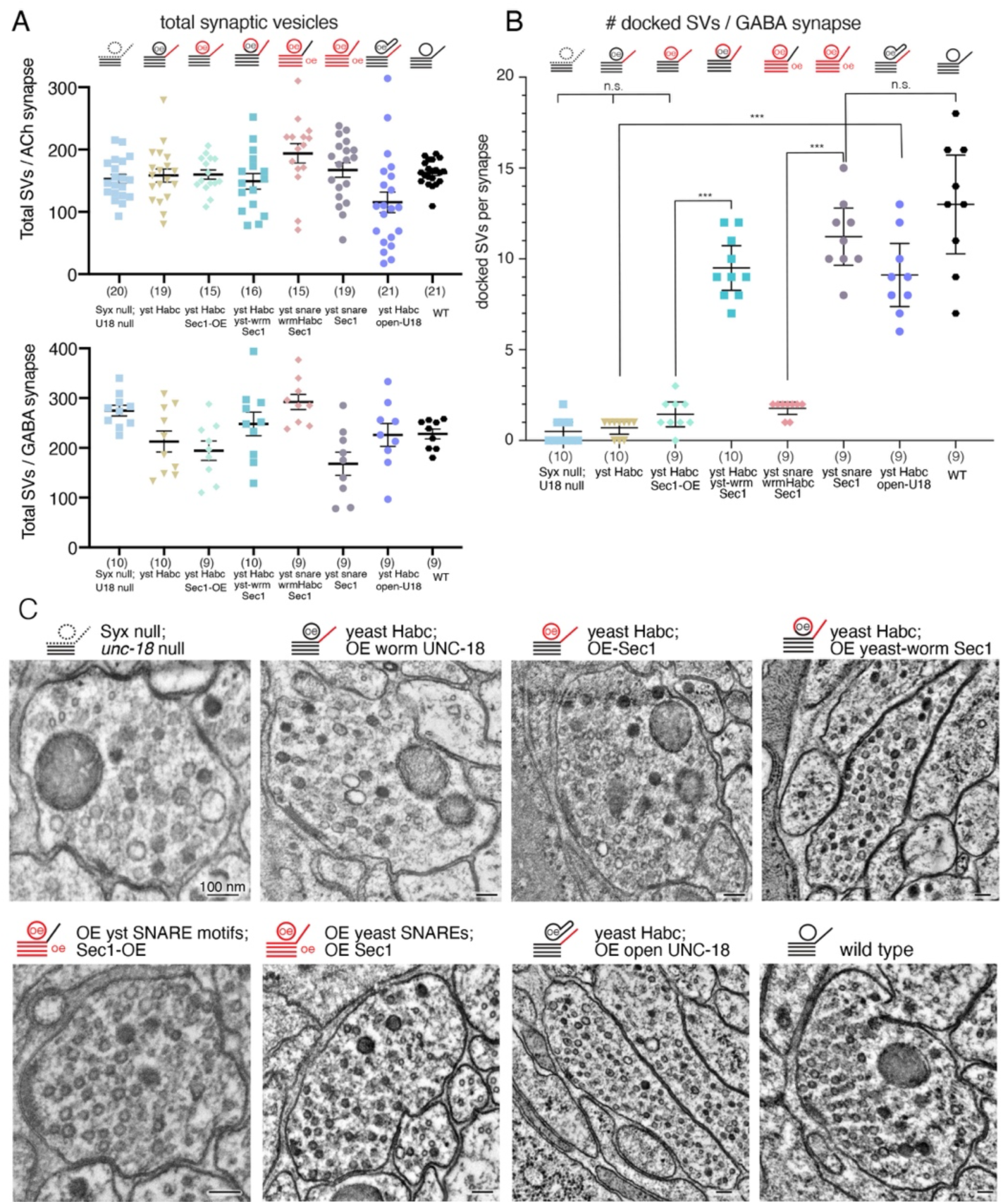
Synaptic vesicle distributions in chimeras. **A,** The total number of vesicles at synapses expressing chimeric syntaxin or SM proteins is not significantly altered. Top: total number of vesicles at ventral nerve cord acetylcholine synapses (VA and VB motor neurons); bottom: total number of vesicles at ventral nerve cord GABA synapses (VD motor neurons). **B,** Synaptic vesicle docking within the active zone is restored when the Habc domain of syntaxin, the SNARE complex, and the SM protein are matched. ‘Open’ UNC-18 bypasses the mismatch of Habc domain and SM protein. Number of docked vesicles in ventral GABA synapses (VD) were quantified for the *syx*-*1 unc-18* double mutant; yeast-Habc chimera overexpressing UNC-18; yeast-Habc chimera overexpressing Sec1p; yeast-Habc chimera overexpressing the Sec1p chimera with ‘groove’ mutations engineered to restore synaptic SNARE interactions; syntaxin mutants overexpressing the full yeast SNARE complex and Sec1p without a matching Habc interaction; syntaxin mutants overexpressing the full yeast SNARE complex and Sec1p with a matching Habc interaction; yeast-Habc chimeras overexpressing worm ‘open’ UNC-18; and wild-type animals. **C,** Representative electron micrographs of the GABA ventral neuromuscular junctions.

**Fig. S5.**
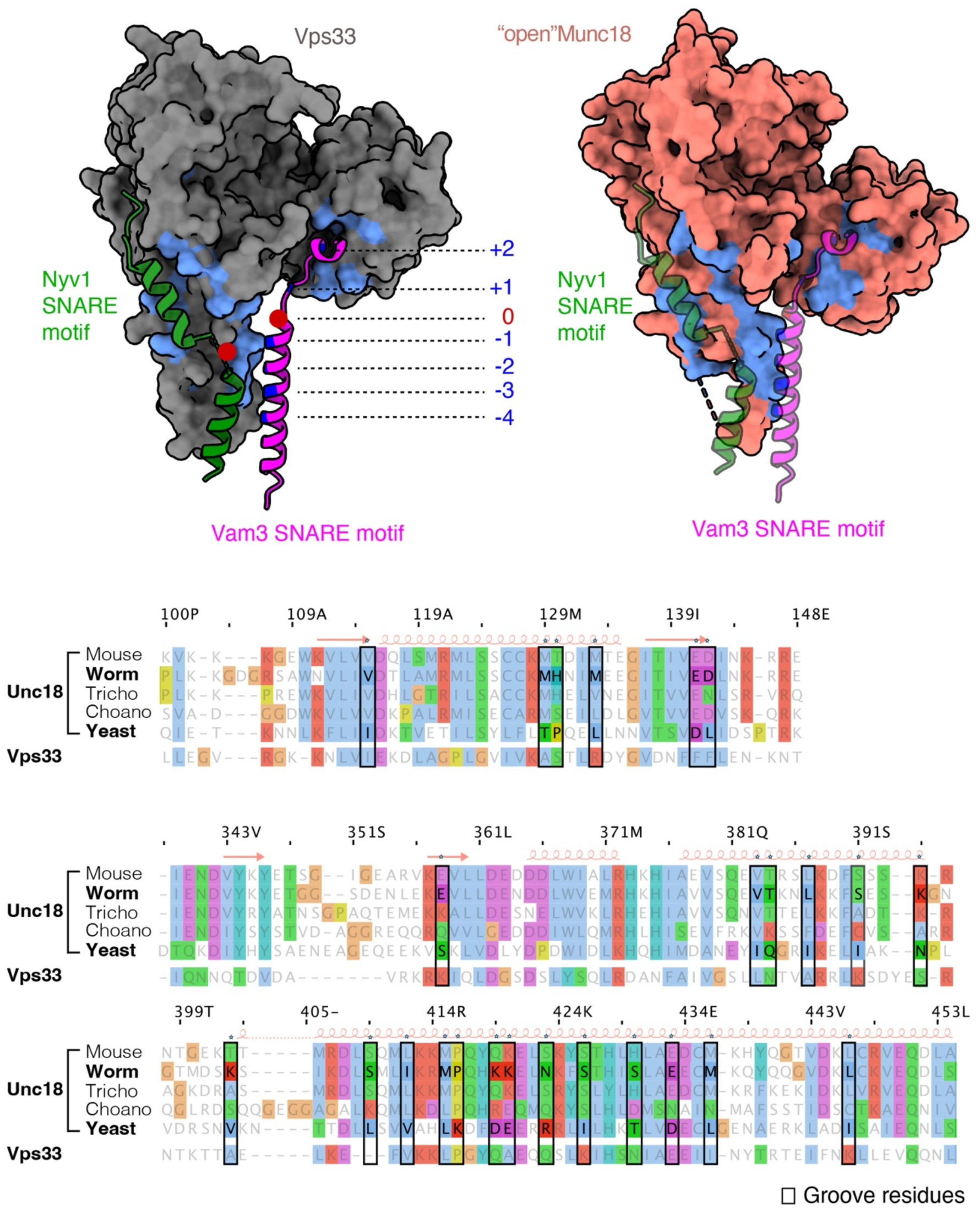
Structure of Vps33 and alignment to SM proteins. The structure of Vps33 bound to its cognate SNAREs served as the guide for engineering residues in Sec1p to interact with the worm SNARE proteins. The Vps33 structure indicates that the Qa-SNARE and the R-SNARE are bound in a partially zippered SNARE complex (PDB codes 5BV0 and 5BUZ). A sequence alignment (below) with the engineered residues boxed indicate the 25 amino acids that were changed in Sec1p to promote interactions with worm SNAREs.

**Fig. S6:**
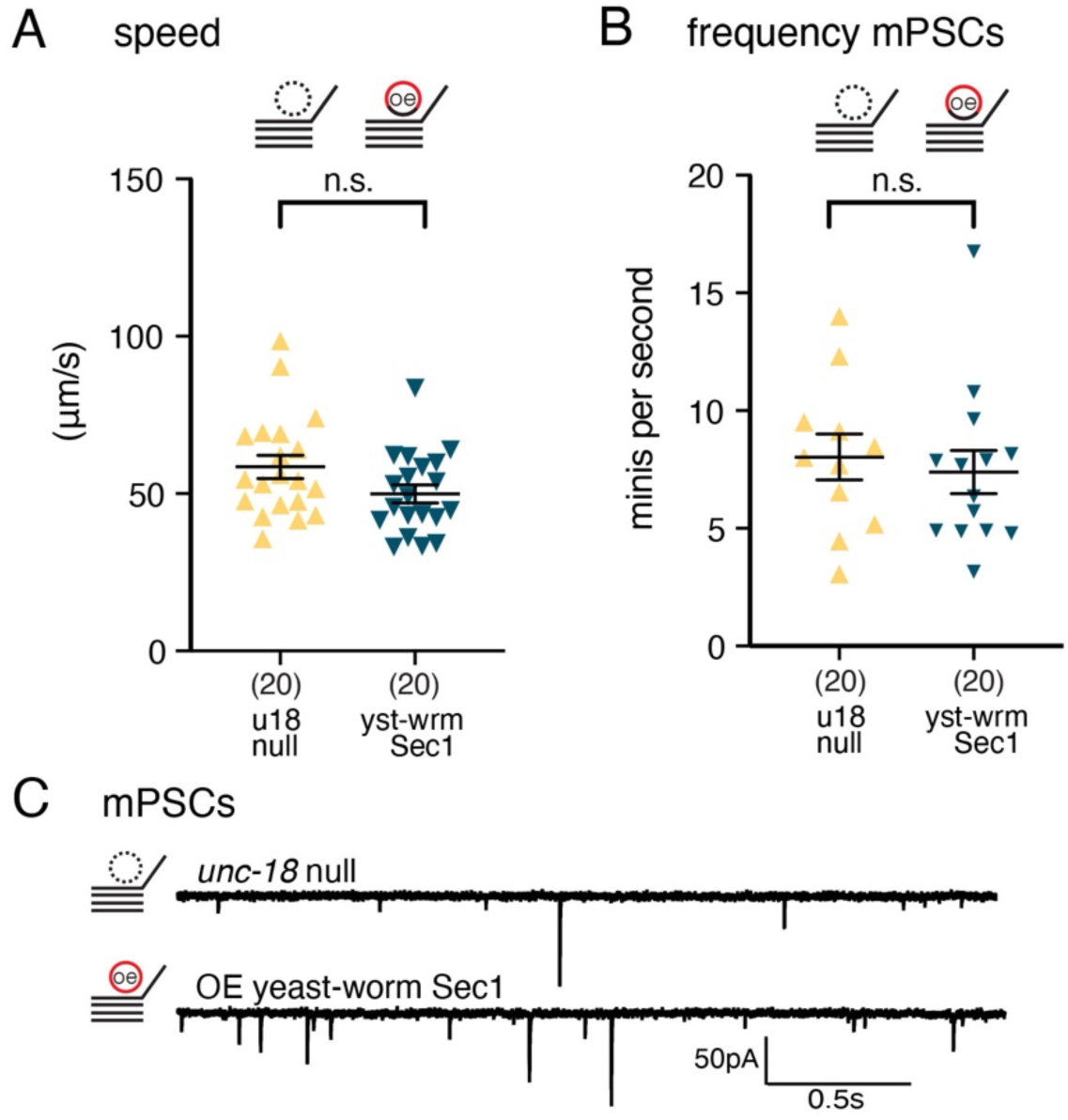
Sec1p chimera is not a gain of function mutant. **A,** Expression of the Sec1p chimera in unc*-18* null animals does not improve locomotion. **B,** The Sec1p chimera does not rescue the frequency of the endogenous miniature postsynaptic currents compared to unc*-18* null animals (*unc-18* null: 8.03 ± 0.98 minis/second, n = 11; *unc-18* null overexpressing the Sec1p chimera: 7.396 ± 0.9159 minis/second, n = 14). Note that the *C. elegans* genome encodes a paralog of *unc-18*, which is expressed ubiquitously, and likely explains the unusually high level of synaptic transmission in *unc-18* null mutants compared to equivalent deletions in other organisms (Cao et al., 2017; Taylor et al., 2021). **C,** Representative traces of endogenous miniature postsynaptic currents (minis) recorded from the body muscle.

**Fig. S7.**
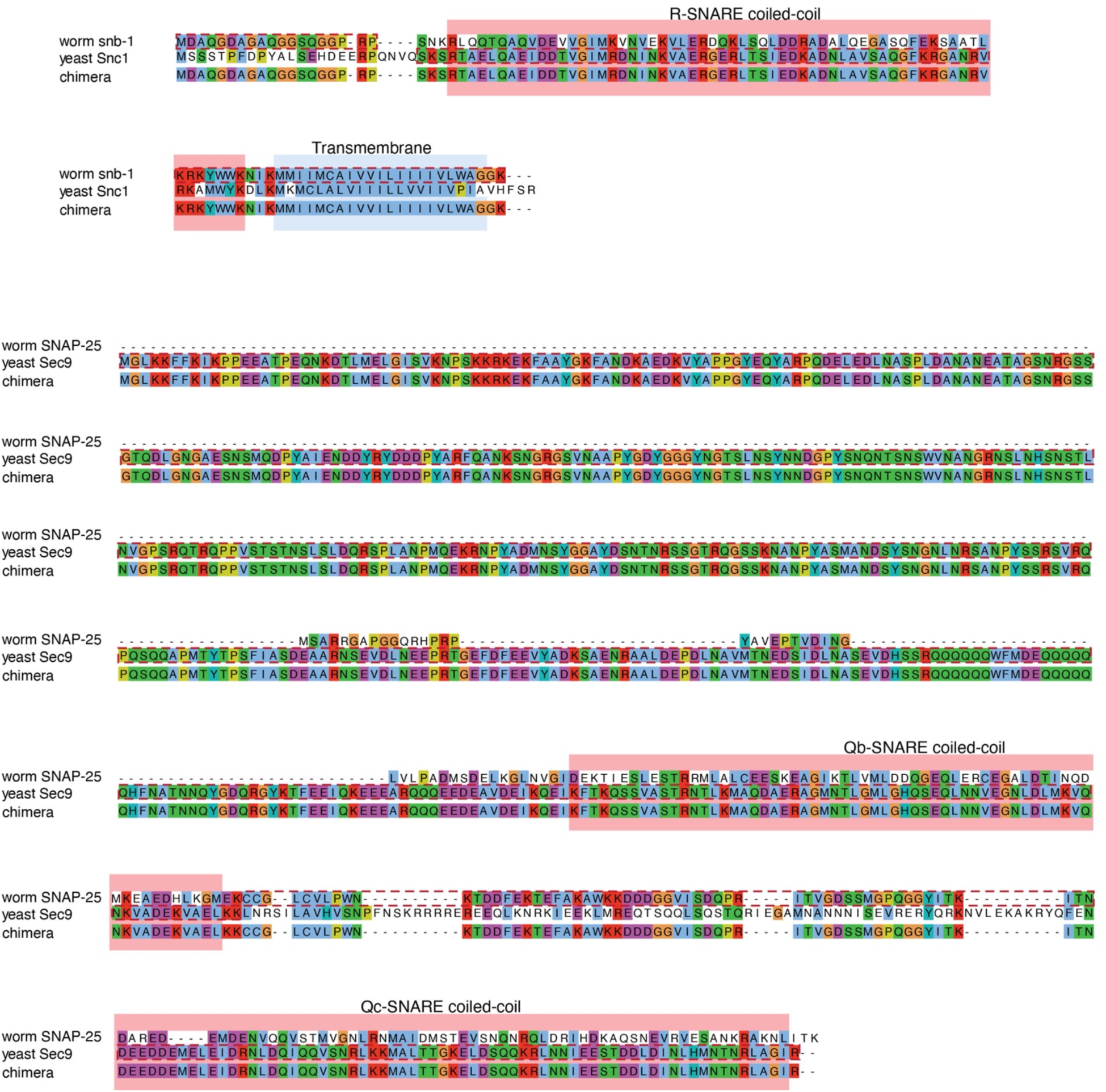
*C. elegans* and yeast SNAREs are highly divergent. Sequence alignments for synaptobrevin and SNAP-25 using worm and yeast sequences are shown. The corresponding yeast-worm chimeras used in this study are shown at the bottom of each alignment. Red dotted boxes indicate the exact region used to build the yeast-worm synaptobrevin and SNAP-25 chimeras. SNARE domains are indicated with a pink box. The alignment uses the default Clustal X color scheme (Larkin et al., 2007).

**Fig. S8.**
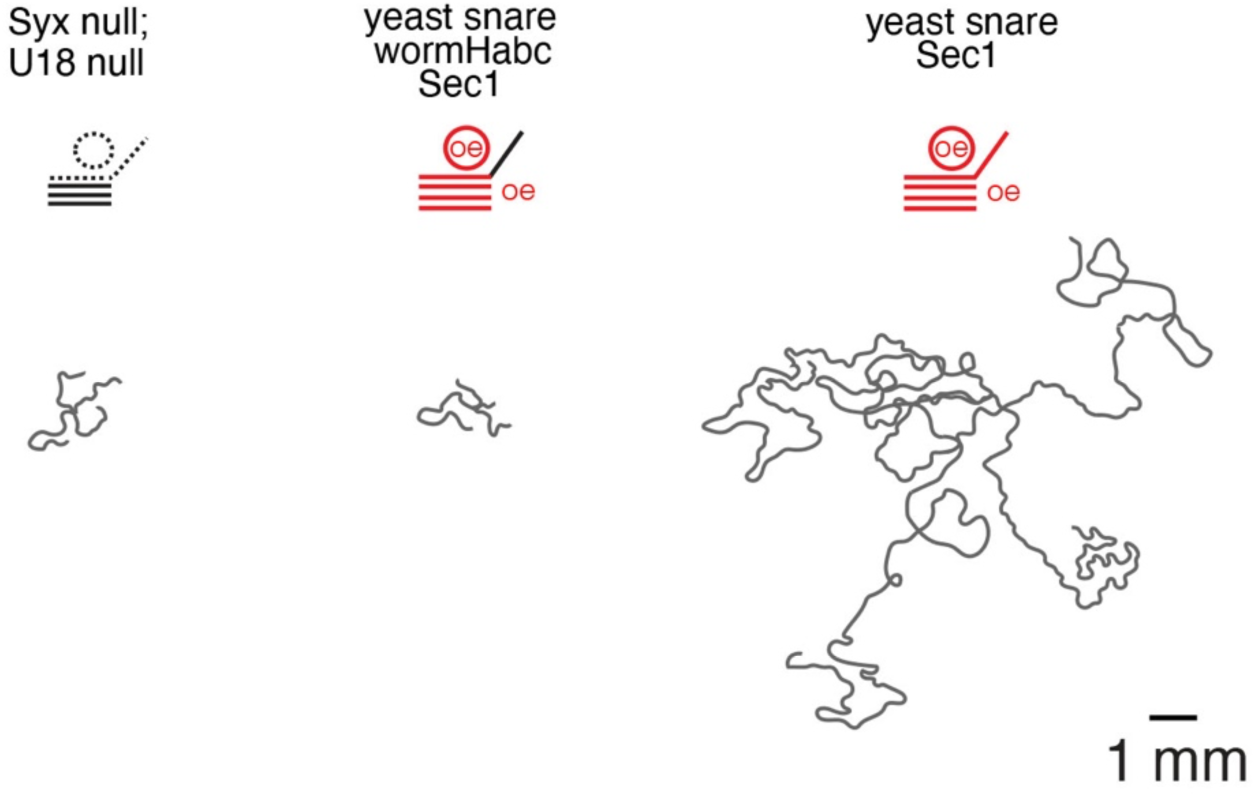
Substitution of the entire yeast SNARE complex along with Sec1p provides significant rescue in *C. elegans*. Representative locomotion trajectories collected over a 5 minute interval, from left to right: *syx*-*1 unc-18* double null mutants; syntaxin mutants overexpressing the full yeast SNARE complex and Sec1p (yeast UNC-18) without a matching Habc interaction; syntaxin mutants overexpressing the full yeast SNARE complex and Sec1p with a matching Habc interaction.

**Table S1.**
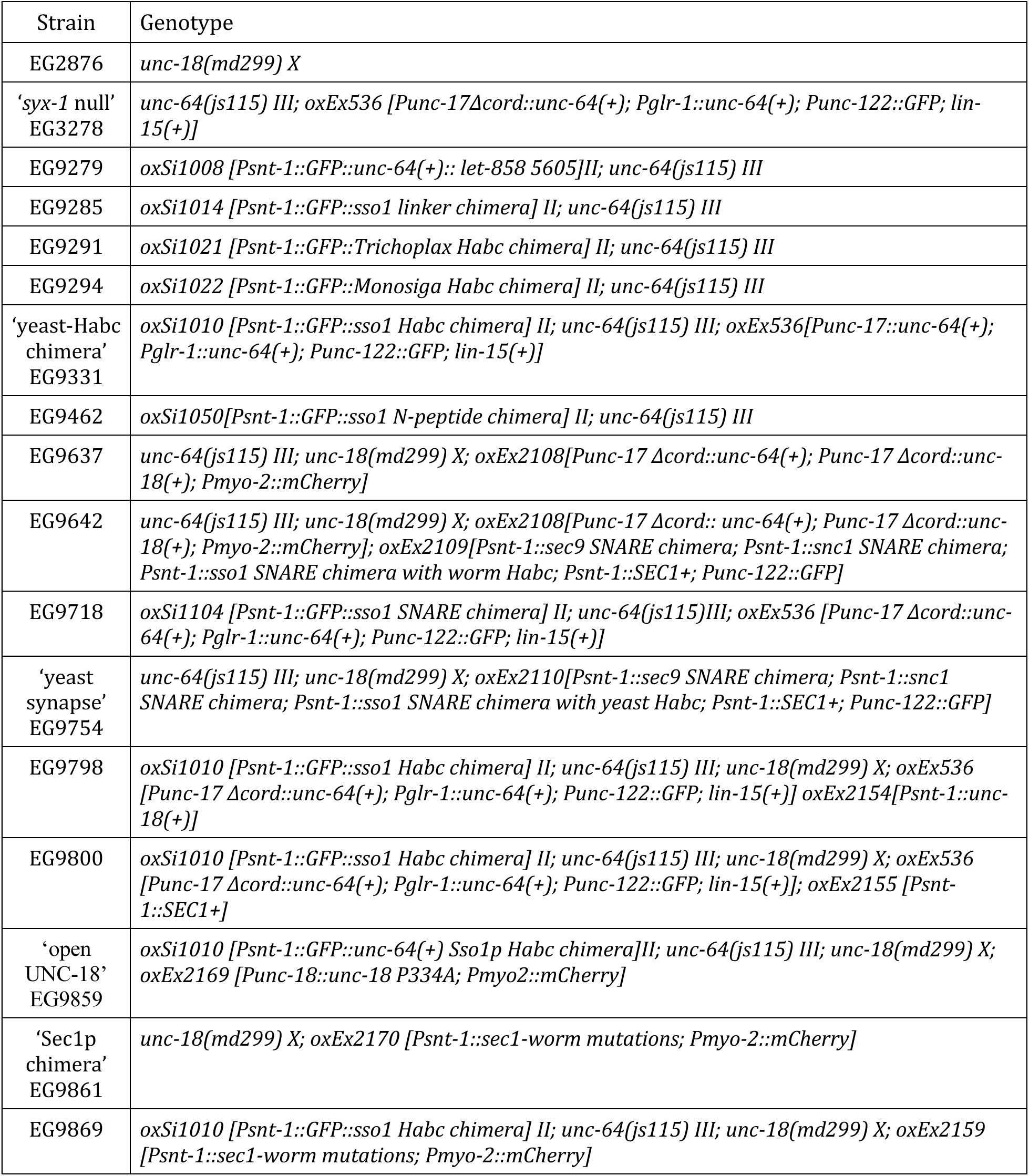
List of strains. Syntaxin is encoded by *unc-64* gene in *C. elegans*, but is also known as *syx-1*. For clarity, we refer to the gene as ‘*syx-1’* in the text, but use the preferred name *unc-64* in the strains list. *unc-64(js115)* is lethal as a homozygote; the ‘*syx-1* null’ strain is rescued by a transgene array expressed in acetylcholine and glutamate neurons in the head; the animal remains paralyzed due to a lack of syntaxin in the ventral nerve cord motor neurons.

